# A single changing hypernetwork to represent (social-)ecological dynamics

**DOI:** 10.1101/2023.10.30.564699

**Authors:** C. Gaucherel, M. Cosme, C. Noûs, F. Pommereau

## Abstract

To understand and manage (social-)ecological systems, we need an intuitive and rigorous way to represent them. Recent ecological studies propose to represent interaction networks into modular graphs, multiplexes and higher-order interactions. Along these lines, we argue here that non-dyadic (non-pairwise) interactions are common in ecology and environmental sciences, necessitating fresh concepts and tools for handling them. In addition, such interaction networks often change sharply, due to appearing and disappearing species and components. We illustrate in a simple example that any ecosystem can be represented by a single hypergraph, here called the ecosystem hypernetwork. Moreover, we highlight that any ecosystem hypernetwork exhibits a changing topology summarizing its long term dynamics (e.g., species extinction/invasion, pollutant or human arrival/migration). Qualitative and discrete-event models developed in computer science appear suitable for modeling hypergraph (topological) dynamics. Hypernetworks thus also provide a conceptual foundation for theoretical as well as more applied studies in ecology (at large), as they form the qualitative backbone of ever-changing ecosystems.

## Introduction

Ecosystems are under threat and we need to think carefully about their representations for many reasons. A sound representation helps to conceptualize the study, to understand and ultimately, to manage ecosystems. Ecological and epistemological works continually propose new representations of these complex objects (Gignoux et al., 2011; Schwartz & Jax, 2011), such as class diagrams and interaction networks (Strogatz, 2001; Loreau et al., 2003; Proulx et al., 2005), to name but two. Such representations should firstly be useful and provide relevant insights. Secondly, even in case of well-understood ecosystems, tracking their changes and managing them efficiently, necessitates dealing with a reliable and intuitive representation. While redefining the ecosystem is outside the scope of this paper, we at least acknowledge that all ecosystems are made up of biotic, abiotic and often anthropogenic components. These components continuously interact through various (bio-ecological, social-economic, physicochemical) processes (e.g., S. Frontier et al., 2008; Gignoux et al., 2011; Gaucherel, 2018). Defined as such, the ecosystem may be, and often is, represented as a set of (material) variables and (abstract) processes (Fontaine et al., 2011; Pilosof et al., 2017; Landi et al., 2018). While a set-based representation does not assign an order to ecosystem content, matrix or graph representations more explicitly specify their complex, directed (oriented) interconnections. Such representations often exhibit specific structures, such as multiple interactions between variables (Sonia Kéfi et al., 2017; Gaucherel & Pommereau, 2019) or nested (Bastolla et al., 2009) and modular sets (Dicks et al., 2002) of variables.

Species networks and social networks provide two textbook cases of network representations. It is common to represent species communities in the form of a graph in which the main variables (species or populations) are represented by nodes and their ecological interactions by edges. Species networks generally consist in sub-systems of an ecosystem, including e.g., food webs, plant-pollinator networks, host-parasite networks and competition networks. Some of these graphical models have also been combined with a dynamic model to examine variations in species abundances and process flows (fluxes) on the graph (Proulx et al., 2005; Delmas et al., 2018; Landi et al., 2018). The use of multilayer, multiplex and multilevel networks has recently been proposed to merge distinct sub-systems in the same network (S Kéfi et al., 2016; Pilosof et al., 2017). These multigraphs are representations combining several initially separate graphs involving the same variables into a multilayered graph.

Latterly, it has been proposed to study *hypergraphs* for their ability to grasp non-dyadic (i.e., non-pairwise) interactions between species, also called *Higher-Order Interactions*, as many ecological interactions appear to be mediated by a third (or more) species (Werner & Peacor, 2003; Golubski et al., 2016; Delmas et al., 2018). Such multi-component interactions thus play an important role in system dynamics. Hypergraphs have rarely been used in biology and ecology (but see Billick & Case, 1994; Klamt et al., 2009 ; Valverde et al., 2020). Several definitions have been proposed for the higher-order interactions they capture: a dyadic interaction modulated by (i.e., in the context of) a third species, a non-additive effect of two species on the per-capita growth of another, or the failure of a null dyadic model (Sanchez, 2019). However, the hypergraphs and higher-order interactions found in the literature are mainly confined to interactions made up of three components and are static. Here, we venture to generalize that i) higher-order interactions not only concern species interactions, ii) are not at all restricted to three-way interactions and, ultimately, iii) often change sharply in the long term. In parallel, social networks are also much used and studied.

In the same vein as species networks, anthropogenic nets can be plotted in graph form, with which humans or human groups shown as nodes and each human-related process (e.g., communication, trading, financial exchange) as an edge connecting the concerned nodes (e.g., Buldyrev et al., 2010; Brummitt et al., 2012). Depending on the question being addressed, such human-related graphs are sometimes purposefully biased by more social, more economic or political processes. Rarely, these interaction networks are merged into the same larger graph for study, such as in decision systems (Tixier et al., 2013; Battiston et al., 2020; Felipe-Lucia et al., 2021). As humans in a society, we understand that social networks are highly dynamic, frequently losing or gaining new components (individuals or groups) and interactions, such as in ecosystem services (Scholes et al., 2013; Mao et al., 2021). In principle, there is nothing to prevent the merging of interaction networks involving human groups with networks involving species once they interact (Felipe-Lucia et al., 2021), and so on with any other (e.g., physicochemical) variable in the same social-ecological system.

In this paper, we will build on the representations above to put forward three interlinked propositions: i) any ecosystem may be efficiently represented by a single graph, here called the *ecosystem network*, whatever the interactions it involves; ii) this single graph would gain from being a hypergraph (called *hypernetwork*), made up of non-dyadic interactions; and iii) the single hypernetwork changes in the long term, hence the need to formally document these changes to improve our understanding of ecosystem dynamics. In other words, we aim to show that the whole (social-)ecological system (i.e., with all the processes and components it involves) can be represented by a single comprehensive graph which may be more accurately defined as a changing hypergraph. In addition, many ecological interactions are non-dyadic, i.e., they involve more than two variables and, reciprocally, most variables are simultaneously involved in many interactions (Billick & Case, 1994; Valverde et al., 2020). In this paper, we will not present new models and new results, rather than exemplify the proposed concepts with our past basic (Gaucherel & Pommereau, 2019; Pommereau et al., 2022a) and applied studies (Mao et al., 2021; Cosme et al., 2022). As a simple illustration, we will model a termite population that is eating, producing mushrooms and reproducing through various processes (Turner, 2009). We will show the numerous advantages of representing the ecosystem as a changing hypernetwork, and will discuss the conceptual, explanatory and practical insights it provides.

## Methods and Results

### A single graph for the ecosystem

To summarize any ecosystem on the basis of its interactions and interaction networks implies an understanding, first and foremost, of what an interaction is. An interaction can be defined in several ways, although most ecologists would accept that it is an abstraction of “real relationships” between species or between any other ecosystem components (Delmas et al., 2018; Landi et al., 2018). For example, a “predation interaction” takes the form of the catching and eating by (at least) one predator individual of (at least) one prey individual (Nakazawa, 2020). Hence, what is usually called predation is an abstract concept, a kind of immaterial process, with a duration in time and aggregating many events. When ecologists study interaction networks to understand or predict their dynamics, they mostly explore quantitative variations in variables (abundances) through various flows (matter and energy exchanges) connecting them (Lobry & Sari, 2015; Nakazawa, 2020) (Fig. 1a).

**Figure 1.**
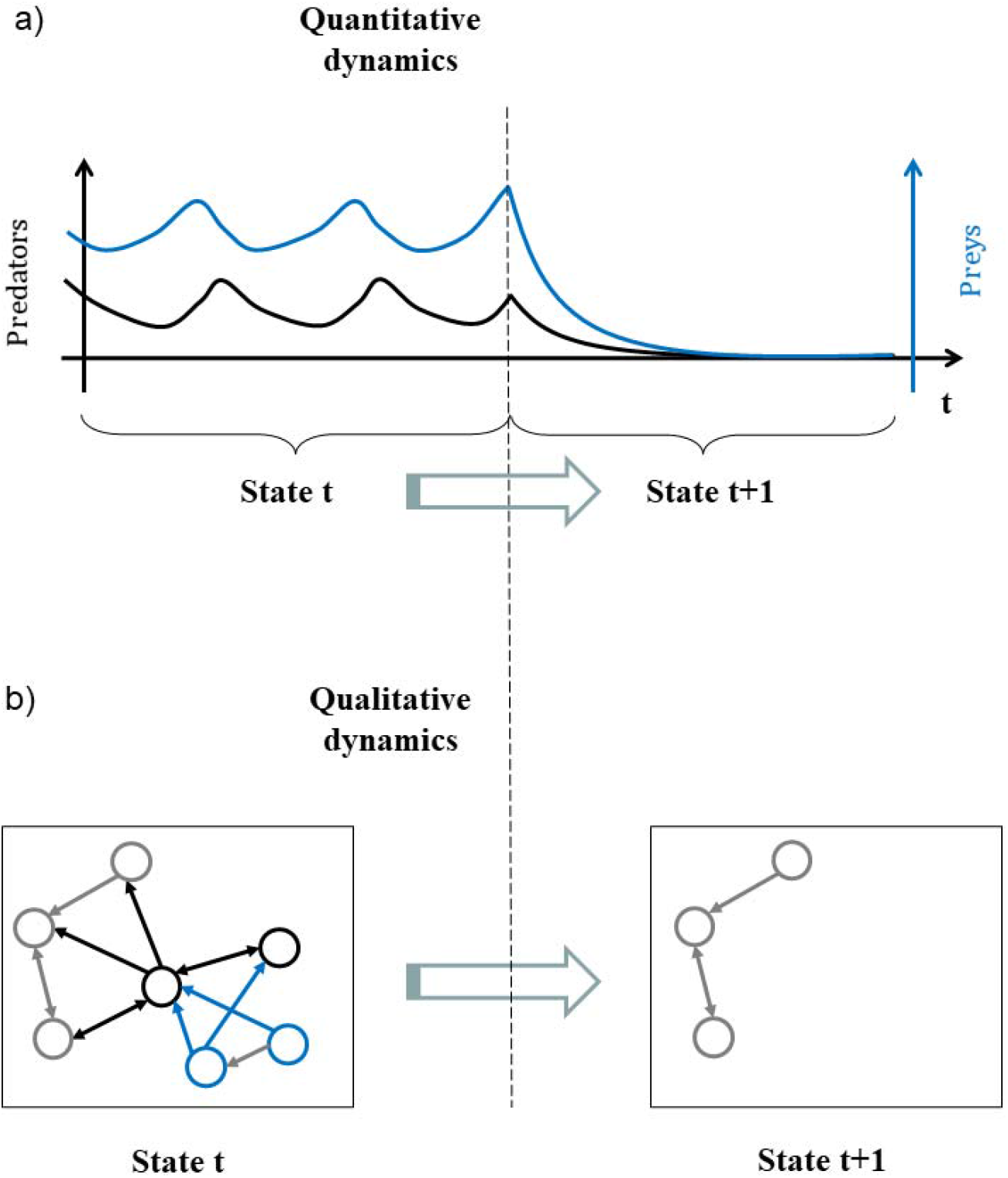
Tentative representations of corresponding quantitative (a) and qualitative (b) predator/prey dynamics. The total biomasses of predator (in black) and prey (blue) populations fluctuate until they collapse (a). In the long term, the trophic network may be represented as a qualitative graph, potentially involving other species (in grey), shifting sharply from one discrete state to the next, lacking the corresponding prey and predator nodes.

Everything that has been said about species may pertain to any other component of a social-ecological system, depending on the question being addressed. For example, the economic interaction stating that “a producer sells their productions to a consumer” is composed of a number of occurrences of isolated and observable sales between two social units (Strogatz, 2001; Gross & Sayama, 2009). Similarly, the social interaction “a scientist communicates their results” is composed of a number of utterances or sentences between the scientist and the audience. The fact that the scientist may simultaneously be communicator, consumer and predator eating a salad for lunch, strongly militates in favor of merging the respective social, economic and ecological networks in which they are involved (Felipe-Lucia et al., 2021). The reverse is also true: many processes require several components to be applied. To “communicate their results”, the scientist may use a computer, the salad they have eaten and potentially studied, and the audience. Hence, as a first step, a single graph can be built in each social-ecological system, depending on the question being addressed (Fig. 1b).

In previous studies, we suggested calling this comprehensive graph the *ecosystem network* (EN) (Gaucherel & Pommereau, 2019; Cosme et al., 2022), as it represents flows and is not simply an abstract mathematical graph. This EN usually concerns a social-ecological system interaction network with no limitations on the natures of its components and processes, insofar as the question being addressed requires all these components. We illustrate here this proposed ecosystem representation with a simplified theoretical termite colony (Table 1). Details of the models can be found in the literature (Gaucherel & Pommereau, 2019; Gaucherel et al., 2020), and minimal information only will be given here for understanding. The termite colony model consists in eight components (graph nodes, Fig. 2a) connected through nine processes (edges), directed from the condition variables to the consequence variables (Table 1). The same graph representing the system may be displayed with several algorithms for highlighting distinct features (Fig. 2a-b), such as ecosystem boundaries or multiscale (nested) structures (Montiglio et al., 2020).

**Table 1.**
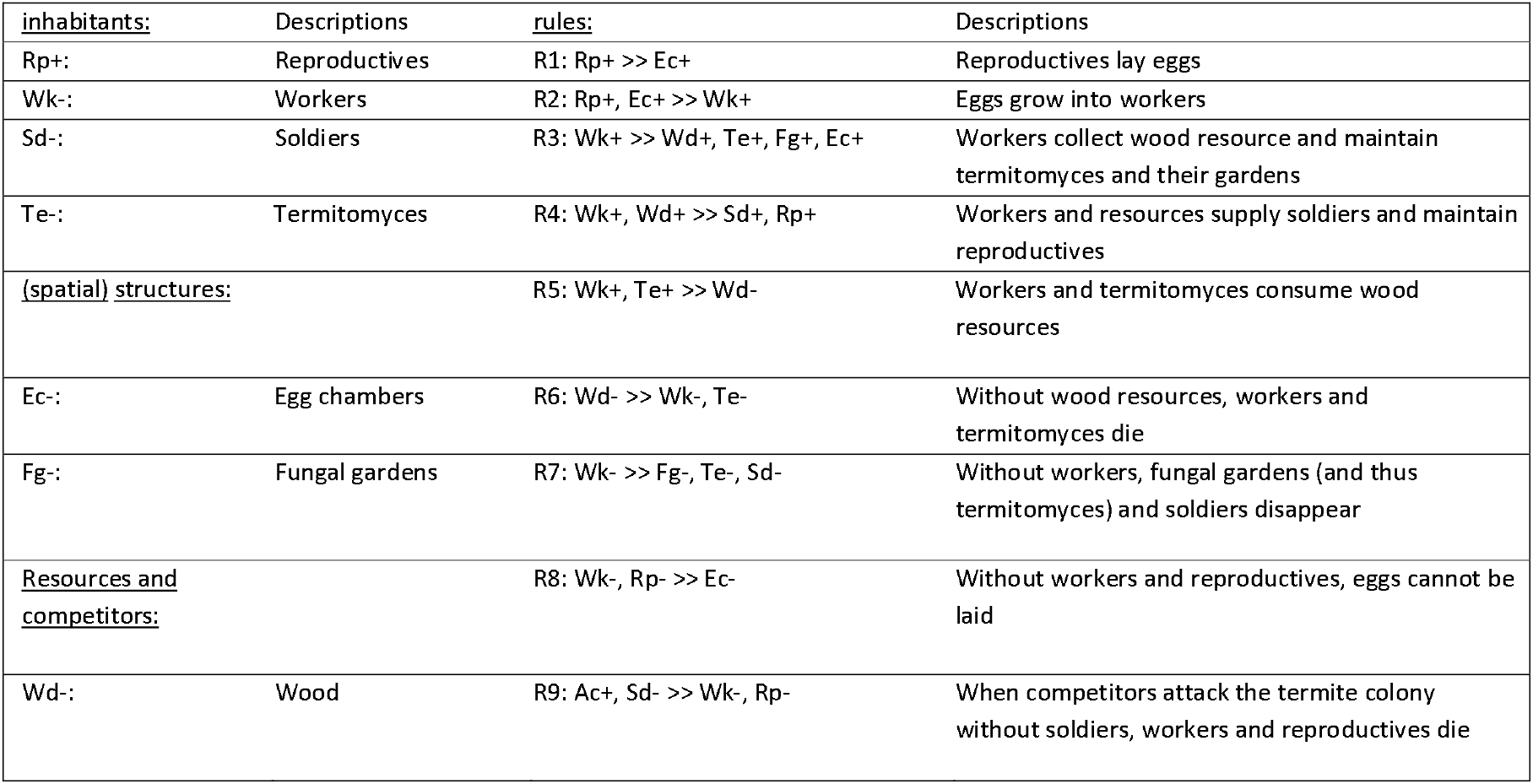
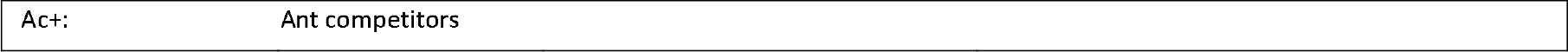
List of the eight termite colony variables with their acronyms, initial states (+ or – beside each variable) and descriptions (columns 1 and 2), and the ten termite colony processes with their descriptions (columns 3 and 4). Simplified from the initial publications (Gaucherel et al., 2017; Gaucherel & Pommereau, 2019; Gaucherel et al., 2020).

**Figure 2.**
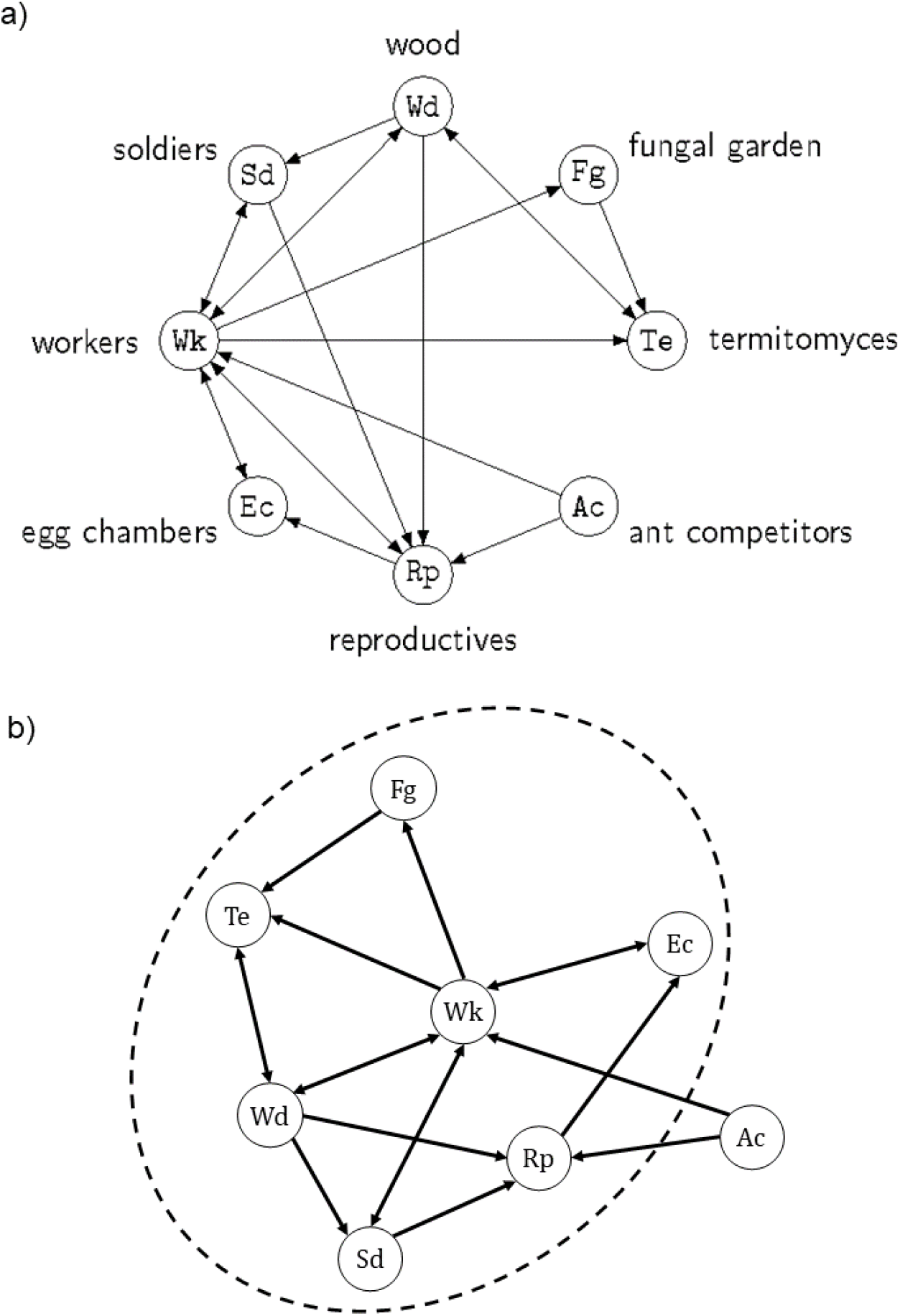
Two different representations of the same network of a termite colony ecosystem (a and b). In the first traditional representation (a), the eight variables (nodes, Table 1) and ten processes (directed edges, Table 1) are represented with no weight or differential location. When a non-circular representation (b) is allowed, here with an appropriate display algorithm (Kamada & Kawai, 1989), certain variables and processes become more central (here, termite workers *Wk*) and others less central (here, competitors *Ac*). The dashed ellipse highlights a possible definition of the ecosystem boundary of this termite ecosystem, on the basis of semi-characterized components (i.e., influencing but not influenced in turn by the system). This representation (b) of the ecosystem is not yet precise.

All the graphs examined so far in various disciplines are indeed “sub-graphs” or modules of the same single “super-graph” of the whole social-ecological system. Often, this is the same node of the EN, representing a material component, which is potentially involved in highly distinct (trophic, physicochemical and/or human-related) interaction networks (Fontaine et al., 2011; Gaucherel, 2019). The reasons that we usually avoid merging all these modular graphs are: specific monodisciplinary questions being addressed; data, knowledge or skills being unavailable; to avoid time-consuming analyses often with limited capabilities; or for all these reasons combined. As seen recently in multilayer, multiplex and multilevel analyses, individual graphs of the same system may be merged into a single larger graph (Fontaine et al., 2011; Pilosof et al., 2017). This observation strongly suggests that each node is connected to nodes of distinct natures, through interactions of various natures too (S Kéfi et al., 2016; Gaucherel et al., 2020). In contrast to these studies, though, we do not make any hypothesis here about the shape of the EN. It may be layered, modular, nested or have another specific structure: we do not need to assume its shape, but instead allow the graph topology (i.e., the neighboring relationships) to emerge from the purported, observed and known interactions (Mao et al., 2021; Cosme et al., 2022).

Furthermore, it follows from this observation that every node, as a component, is a super-graph (made up of finer sub-nodes) in itself. Indeed, it is always possible, and sometimes relevant, to subdivide a population node into individuals or groups (e.g., population stages, age-wise, sex-wise), and to emphasize their various actions, at various levels of organization. All these representations need to be useful for the question being addressed, though, as it is often pointless to subdivide a component into a new subgraph to achieve a more detailed representation of the system. This remains valid for a specific interaction which it is pointless to subdivide into several (dyadic) processes. Whenever it is relevant for the question being addressed to zoom in (to “unfold”) and/or zoom out (“fold”) part of the single graph under examination, it is possible to split nodes into sub-nodes connected to other nodes already present with distinct processes (Gaucherel, 2019; Montiglio et al., 2020). Such a procedure obviously requires caution as it quickly encompasses distinct levels of organization (or scales) too.

### From graph to hypergraph

In real systems, most variables are connected to many others, such as the aforementioned scientist who interacts with other scientists for the purpose of communication, with salads as food and with producers for shopping, through the processes of talking, eating and buying. Conversely, most processes involve many components (variables) simultaneously, such as the communication process requiring the scientist, potentially an organism under examination (the salad), and the audience. This observation remains valid for any subgraph and discipline such as, for example, a physicochemical graph of chemical reactions requiring reactants and catalyzers (Fontana & Buss, 1994; Cumming et al., 2014; Fages et al., 2018). For these reasons, most interactions are called non-dyadic (i.e., not reduced to dyadic interactions), a point that has been made recently for ecological networks too (Werner & Peacor, 2003; Golubski et al., 2016; Delmas et al., 2018). In ecology and elsewhere, it may be that such non-dyadic interactions are the norm rather than the exception. This leads to complex ENs with a large number of complex interactions, which require appropriate concepts for disentangling them. Computer science has developed conceptual and technical tools to handle such non-dyadic interactions and compute their dynamic consequences. Known as *hypergraphs*, a number of associated tools have been developed to handle them (e.g., Reisig, 2013; Battiston et al., 2020), although they remain somewhat limited, especially for non-specialist practitioners, as compared to traditional dyadic graphs.

A *hypergraph* is a generalization of a graph in which edges, called *hyperedges*, may have several incoming and outgoing nodes. In ecology, any variable may be connected through many processes and, above all, any process may connect many variables, as in the case of the termite colony (Table 1). In our studies, we use hypergraphs by dictating that nodes are the material components of social-ecological systems and that edges are the abstract processes involving these components (e.g., Mao et al., 2021; Cosme et al., 2022). Hence, any hypergraph is a means of representing the state of the ecosystem via the composition of its components, while still considering their associated interactions. In addition, it is possible to define the processes involved on the basis of equations or rules that represent the components’ interactions. Such process definitions, be they dyadic or not, naturally direct the associated EN hyperedges in most environmental studies. Any hypergraph may be represented by a Petri net, among other convenient tools developed in computer science (Fig. 3b). Petri nets were developed in the 1930s for modeling chemical reactions requiring precisely such connections and have since been used in many other fields (Pommereau, 2010; Reisig, 2013; Pommereau et al., 2022b). In such a representation, variables are known as *places* (round-shaped nodes) and interact through processes called transitions (square-shaped nodes), shown as arcs. Any hyperedge may be represented by the set of arcs (or *hyperarcs*) of a Petri net, connecting the nodes as the pre-condition and other nodes as the post-condition of the focal transition (Fig. 3b). Petri net arcs may be weighted, and places may be multi-valued, holding varied numbers of tokens, thus constraining their flows in the Petri net. Such bipartite graphs are equivalent to hypergraphs as they allow an arbitrary number of nodes to be connected to an arbitrary number of edges and vice versa (e.g., the termite colony EN, Fig. 3c).

**Figure 3.**
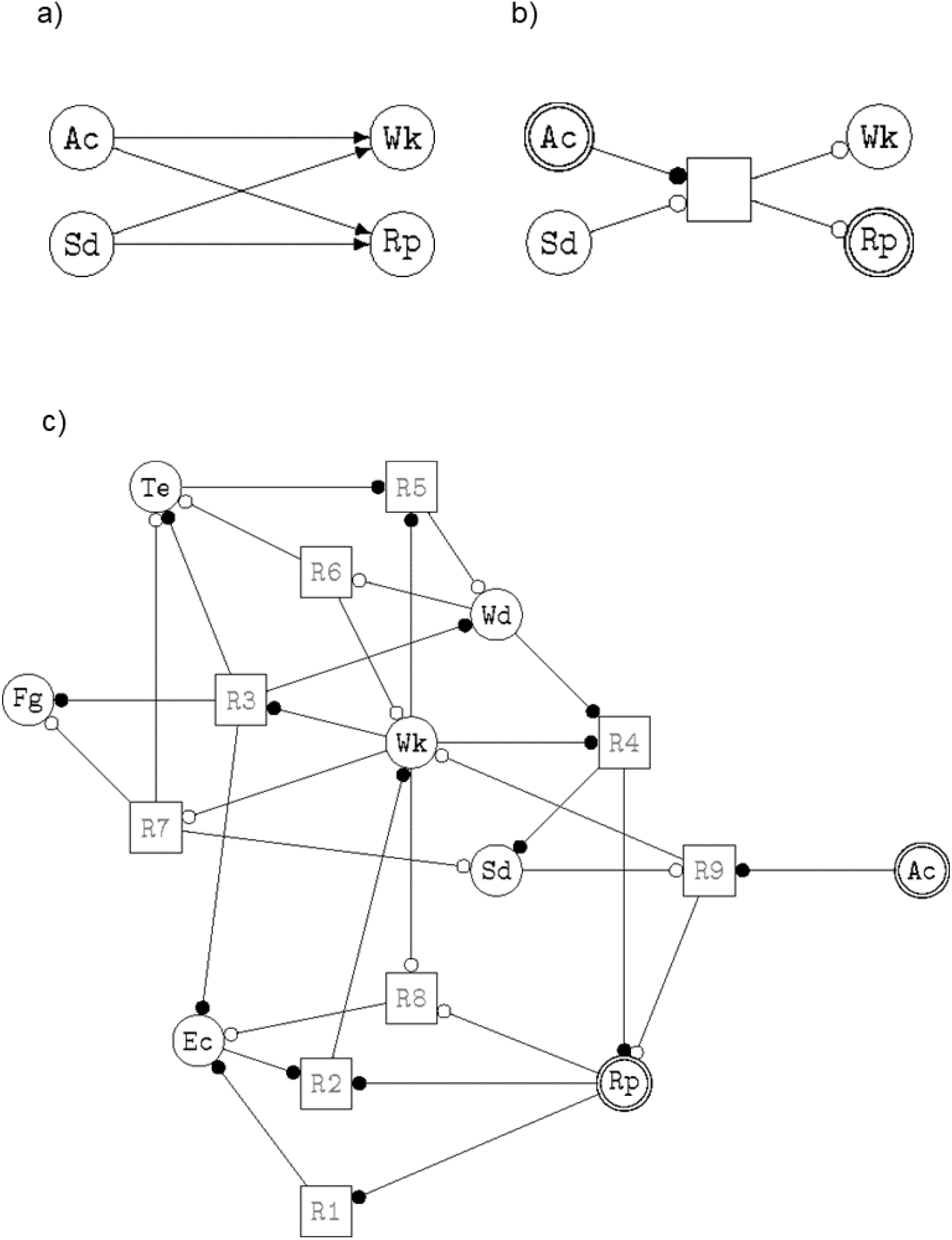
Illustrations of the symbols chosen here to accurately represent the ecosystem network hypergraph (a) and the full representation of the termite hypergraph (b, the same ecosystem as in Fig. 1). The sub-hypergraph corresponding to the last rule (R9) of the termite ecosystem (Table 1) is displayed: *Ac+, Sd->> Wk-, Rp-* with its corresponding traditional graph (a) and our proposed hypergraph notation (b). See the main text and (Pommereau et al., 2022a, 2022b) for explanations of each symbol indicating how tokens should circulate in the hypernetwork. The full ecosystem network hypergraph of the termite colony is displayed (c) with the corresponding symbols (b), and with the variable and process names found in Table 1.

Shifting to the termite hypergraph, we define specific symbols to signify the Boolean nature of each node (component) in the rule (process) (Fig. 3a-b). Circular nodes are variables and represent social-ecological components. A node is drawn with double lines if it is initially present in the system (Boolean value +), or a single line otherwise (-). When a variable is present, its node has a token (not shown), otherwise it does not. Square-shaped nodes are transitions and represent social-ecological processes. Each process is defined graphically by a condition, arcs (round-tipped arrows, Fig. 3b), a transition node and a consequence, and forms a directed hyperedge. A transition node “reads” and “writes” the values (tokens) of variables. When an incoming arc (from a variable node to a transition node) has a black (resp. white) dot, then the transition node “expects” the variable to have a token, i.e., to be present (resp. absent). Conversely, when an outgoing arc (from transition node to variable) has a black (resp. white) dot, then the action activates (resp. inactivates) the variable. In the Petri net, a variable activation (resp. inactivation) corresponds to token production (resp. consumption). A variable may be present in the condition and in the consequence of the same action. In this case, there will be a dot at each end of the arc connecting the action to the variable. The resulting graph (Fig. 3c) unambiguously represents the termite ecosystem EN, and thus explicitly links network structure and dynamics.

An example of non-dyadic interaction may be found in the last process (R9): Ac+, Sd- >> Wk-, Rp-; stating that termite workers (Wk) and reproductives (Rp) may both die when competitors (Ac) attack the termite colony in the absence of soldiers (Sd) to defend it. Using the model’s terminology, the condition “presence of competitors and absence of soldiers” enables and triggers the consequence “absence of workers and reproductives”. In this case, a traditional graph representation (Fig. 3a) is not appropriate, as it hides the exact connections of this four-node interaction. To be rigorous and to retain all the ecosystemic information, we need a graph that explicitly shows which variables interact with each other, and how. Inspired by Petri net notations, we propose to replace hyper-arcs (initially connecting nodes) with another node type (like transitions in Petri nets, Fig. 3b). The resulting graph (Fig. 3c) unambiguously represents the termite ecosystem EN (Fig. 2b), and thus explicitly links network structure and its potential dynamics.

Simultaneously, it becomes relevant to analyze the EN structure itself and to highlight the graph topology for ecological insights (Fig. 2b-3b). Here, no shape is assumed for the EN; instead, the graph topology is allowed to emerge from the chosen variables and known interactions, bringing interacting variables closer and moving non-interacting variables further out. With an appropriate graph layout (Fig.2b), algorithms and graph analyses (Kamada & Kawai, 1989), it is possible to produce representations that help in understanding node properties (e.g., whether they influence or are influenced by other nodes) and many other properties (e.g., betweenness, connectivity index) as is already routinely the case in ecology (e.g., Urban & Keitt, 2001; Foltête & Giraudoux, 2012). For example, a relevant *boundary* of the ecosystem under examination may be computed by excluding nodes that influence others but are not influenced in turn (e.g., the sun and other so-called *control variables*), and including all the others (e.g., a species simultaneously eating and being eaten) (dashed ellipse, Fig. 2b).

### Qualitative ecosystem dynamics

Graphs and hypergraphs are by definition discrete. However, the interaction network they represent might not be (Dicks et al., 2002; Landi et al., 2018; Valverde et al., 2020), as their nodes may represent continuous variables and their edges continuous flows between them (Fig. 1a). In this paper, we propose to approximate the interaction network of any ecosystem with a qualitative hypergraph (still generically denoted as ‘EN’ hereafter). This hypergraph carries information about the potential qualitative dynamics of components, namely their appearance and disappearance (Fig. 1b). This description is relevant for long-term dynamics, as an observer who waits long enough will no longer measure flows and abundances, but appearance and disappearance of system components (and processes). In this spirit, we define a system component as a *Boolean* variable, i.e. having two discrete values, describing the functional presence/absence of the component in the system. A component that is functionally present is considered to pass a predefined threshold, present in most non-linear processes (R. Thomas, 1973), and to be able to influence (or be influenced by and possibly, no more influence) other components (Fig. 1b). Conversely, a process (potentially generating an *event*) is defined as a discrete change in the values of certain variables (the consequence) triggered by other variables (the condition). According to Petri net formalism (Pommereau, 2010; Pommereau et al., 2022b), these events are represented by *rules* written as “condition >> consequence” (Table 1). The change in one (or more) variable(s) is called a transition, which creates a new system state.

The sequences of states and transitions are the system *dynamics*. The dynamics of a qualitative EN would enable the sharp changes in the long-term dynamics of the ecosystem to be understood. In a way, the single qualitative EN constitutes the *skeleton*, a kind of backbone of the (social-)ecological system under study, intentionally neglects subtle variations and flows (Gaucherel & Pommereau, 2019; Cosme et al., 2022), i.e., the component dynamics, and enables the computing of qualitative ecosystem dynamics, the network dynamics (Gaucherel et al., 2017; Gaucherel, 2019). An EN hosts the tangled dynamics of species abundances and other components and of flows between them. Many studies have developed powerful models to simulate and understand such dynamics in networks with frozen topology (Thébault & Fontaine, 2010; S. Kéfi et al., 2015). Using discrete-event models, it becomes possible to model changing networks and their discrete dynamics (i.e., state and transitions). A system state is defined by the values of all the variables. In the graph, a system state corresponds to a given token distribution (called the *marking* in Petri nets) among the nodes (Fig 4a). A state transition occurs when the marking changes, that is to say, when one or more tokens are produced and/or consumed in the graph.

**Figure 4.**
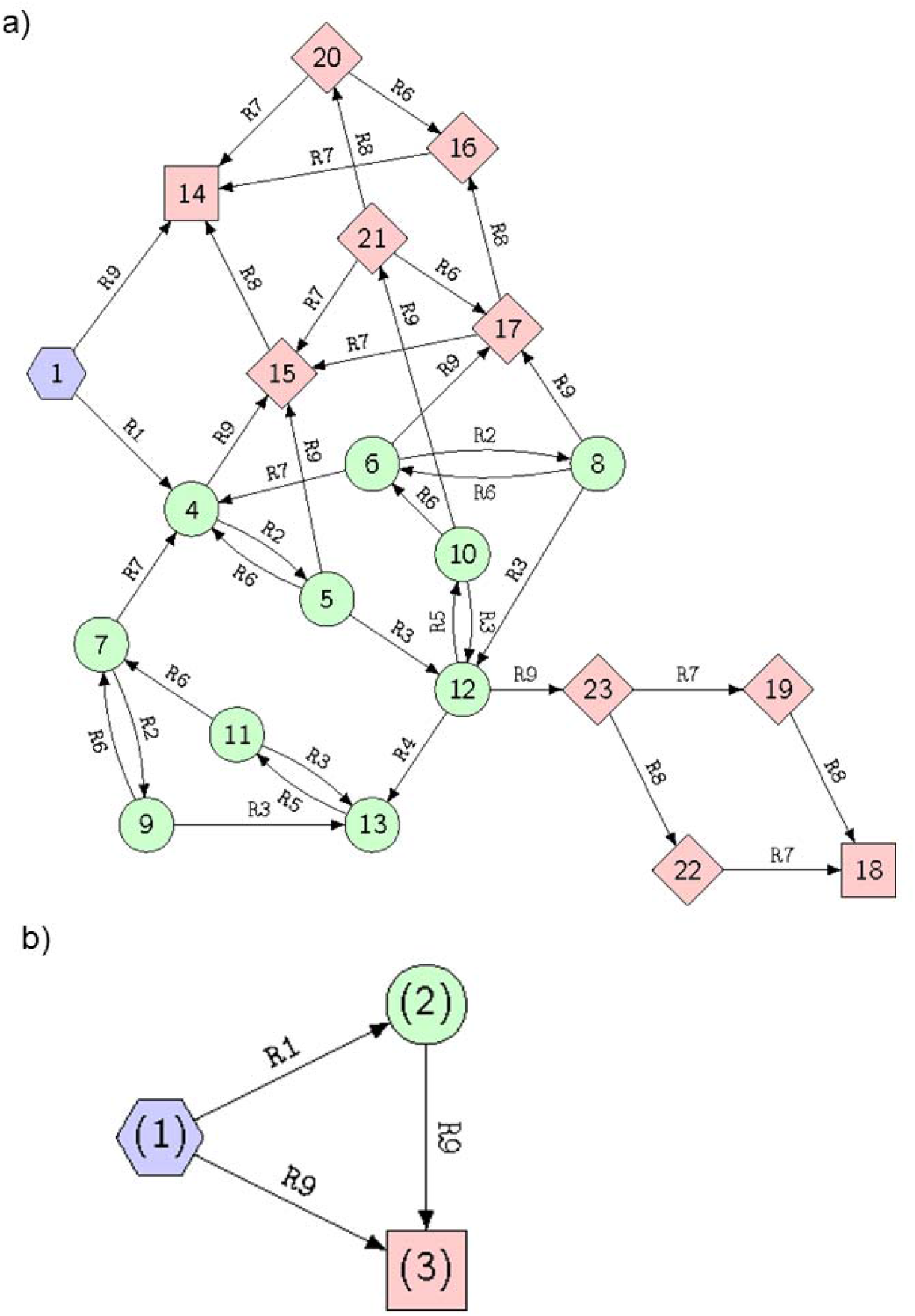
The termite ecosystem state spaces (or STGs), full (a) and merged (b), i.e., the dynamics computed from its corresponding ecosystem network hypergraph (Gaucherel & Pommereau, 2019; Pommereau et al., 2022a). The pathways start from the initial state (hexagon), follow various states according to process rules (defined in Table 1), cycle into sets of states (colors), and finally reach deadlocks (squares). The node labels are arbitrary identifiers of each state (a) and of each structural stability (in brackets, b), while each edge is labeled by the rule (Table 1) responsible for the system change.

Starting from an initial state (blue hexagon in Fig 4a, and column 1 in Table 1), the graph enables the computation of all reachable system states and transitions (following green and pink states). The modelled system may then bifurcate in state n°4 by following rule R1 (“Reproductives lay eggs”, Table 1) or in state n°14 by following rule R9 (“workers and reproductives die”), and so on. These states and transitions are usually represented as a State-Transition Graph (STG, Fig. 4a) (Berthomieu et al., 2004; Reisig, 2013). In this STG, any cycle (i.e., a set of mutually reachable states, also called structural stabilities) or fixed state (a state with no outgoing transition, also called deadlocks) becomes now identifiable and informative (Gaucherel & Pommereau, 2019; Gaucherel et al., 2020) (colors in Fig. 4a). Hence, it is possible to automatically compute the discrete-event dynamics of the system from the hypergraph structure (Fig 4b).

This is precisely what we modeled in several studies from a chosen EN of temperate and tropical social-ecological systems (Mao et al., 2021; Cosme et al., 2022). By using specific tools, it is also possible to formalize specific questions about the dynamics displayed in the STG, without explicitly computing them, a relevant advantage when it becomes too large to be fully computed (Fages et al., 2004; Largouët et al., 2012; C. Thomas et al., 2022). In addition to an intelligible visualization of ecosystem structure and dynamics, we now have the benefit of the whole toolbox associated with such representations (Fig. 4b), as with the Petri nets that computer science has developed (Berthomieu et al., 2004; Rodríguez & Schwoon, 2013; Pommereau et al., 2022b). The demonstration of the relevance and application of Petri nets and related concepts is outside the scope of this conceptual paper, but interested readers may refer to recent papers (e.g., Gaucherel & Pommereau, 2019; Mao et al., 2021; Cosme et al., 2022). Such models automatically produce STGs comparable to those already developed in many empirical ecological studies (Stringham et al., 2003; Bestelmeyer et al., 2017). Looking ahead, several computing tools are even able to automatically detect various kinds of topological structures and regularities/invariants directly within the EN (Rodríguez & Schwoon, 2013; C. Thomas et al., 2022). For example, in theory, individuals or populations that may exhibit cycles or that may collapse could be detected by such invariants directly within the EN representation (Fig. 4b). This view supports an innovative conceptual representation of the ecosystem.

### A changing (and spatial) hypergraph

Disregarding for a moment variable abundances and process flows, this EN is in no way frozen. In the long term, each component and each process potentially disappears and (re)appears in line with the presence/absence of its material components (Fig. 1b). They can enter (e.g., species invasions or pollutants) and/or leave (species extinctions or migrations) the system being studied. Our aim is to model network dynamics, i.e., a changing topology, not to be confused with the dynamics of components in a static network (i.e., flows “carried” by the frozen graph). In the long term, the nodes and edges of the graphical representation themselves become fragile and ephemeral, depending on the other connected variables, what previous studies have called *ecosystem development* (Gaucherel et al., 2017; Gaucherel, 2018). Conversely, in short-term dynamics with a similar level of detail, the EN retains the same structure (Fig. 1 or any state in Fig 4a). This assumption of a fixed topology is still widely used in most ecological studies today (e.g., Proulx et al., 2005; Golubski et al., 2016; S Kéfi et al., 2016). We posit that such topological changes, or ecosystem development, strongly differ in nature to ecosystem *functioning*. We developed a number of models and tools to analyze such qualitative dynamics. In the termite colony toy-model, the STG computed from the chosen initial state is well suited to detecting stabilities, collapses, and tipping points (Fig. 4b) in ecosystem pathways (Gaucherel et al., 2020), if any. Today, we are applying this framework to large and more realistic observed social-ecological systems (e.g., Cosme et al., 2022; Cosme et al., 2023), sometimes made up of dozens of components and hundreds of interactions of contrasting nature.

In the long term, the (qualitative) EN topology is continuously changing, and these structural (or network) dynamics may be largely disconnected from short-term component dynamics. Any sharp change in the system will change the network topology, which is modeled by the token flow. Modeling such a changing topology is highly difficult, however, because it implies anticipating any disappearance and (above all) appearance of components. Practically, it is easier to initially assume the *maximal* EN topology by including all possible components, and then mimicking their possible disappearance by removing their associated nodes and/or tokens. For example, in our past studies, we model structural changes in the EN representing the appearance and disappearance of components by changing the Boolean values of their associated variables and nodes (Fig. 1b). This is known as the EDEN (for Ecological and Social Discrete-Event Network) framework (Gaucherel & Pommereau, 2019; Pommereau et al., 2022a). Such changes are generated by certain rules representing system processes and their related transition events. The STG precisely computes the successive topologies of the EN, considering the present and absent components and processes of the termite colony (nodes in Fig. 4a). Hence, any STG state displays a distinct EN topology, mimicked by the variable changing states (Fig. 3). This makes the EN an ever-changing hypergraph whose topology needs to be reconstructed for an understanding of the long-term dynamics.

The EN representation does not fit with the traditional interaction networks in ecology or environmental sciences as, firstly, we proposed to collect all the interaction networks present in an ecosystem via qualitative approximation (Fig. 2); and secondly, as we proposed rigorous conventions for this representation (Fig. 3), inspired by Petri nets which are well adapted for handling such mathematical objects (Pommereau, 2010; Rodríguez & Schwoon, 2013). It is striking that most models used in ecology today (mainly ordinary differential equation systems) are not appropriate for handling dynamic systems on dynamic structures (called DS2, Giavitto, 2003). Here, the central idea was to integrate all relevant interactions for the question being addressed into the same graph, and to study the coarse (topological) graph dynamics. The EN is a coherent and convenient concept for summarizing all ecosystem components, whatever their nature and complex interactions. It provides a way to achieve the often sought and rarely attained integration of social-ecological systems (Ostrom, 2009; Mao et al., 2021). Another advantage of qualitative ENs is their intelligibility (Dambacher et al., 2003; Thébault & Fontaine, 2010), even at larger sizes, while quantitative models generally require a numerical analysis and act like black boxes. It is possible to define extensions of the notation presented above that would use as many tokens as required to model quantitative variations for each (multi-valued) variable, but at the cost of an explosion in STG sizes and in the model’s computation. And nothing prevents, then, the transformation of hypergraphs into hybrid models combined with systems of differential equations for more quantitative analysis (Samaga & Klamt, 2013). While most traditional models in ecology and environmental sciences are limited to a dozen or so variables, with numerical computations, our qualitative models often handle dozens of variables (Cosme et al., 2023).

In addition, the EN appears heuristic in that, in common with traditional interaction networks, it reveals the ecosystem’s internal structure: the EN topology collects internal variables and their specific interactions, while highlighting external variables too (Fig. 2b, dashed ellipse). Such external nodes are control variables that influence (toward ecosystem centrality) without being influenced in turn. In other words, the boundaries of the ecosystem or the “skin”, become visible on the skeleton. Nonetheless, i) the ecosystem boundary is not necessarily a spatial or specific (e.g., species-centered) boundary, rather than a multidimensional functional boundary (Fig. 2b), and ii) the hypergraph representation is not hampered by the boundary definition so often debated (Gignoux et al., 2011; Gaucherel, 2018). It naturally emerges here from the processes listed in the model, and from the question being addressed. When conceived in this way, it becomes obvious that the ecosystem does not conserve matter and energy, and is thermodynamically open (Jorgensen, 2001; S. P.-V. Frontier, D. et al., 2008). Conversely, ecosystem variables located in the center of the EN hypergraph (e.g., termite workers in Fig. 2b), interact with more variables in more diversified ways. They do not play a higher role, but they are more closely related to the question being addressed, as the modeler explicitly considers its possible impacts and influences on the rest of the system. Yet, we warn the reader that it may be dangerous to formulate potentially naive interpretations on the basis of a displayed structure only.

In parallel, providing another perspective, all the points discussed previously may be included in spatially implicit or explicit ecosystem models. Indeed, space itself is easily represented as a graph: in the case of a spatially explicit ecosystem, the localities involved are potentially represented as nodes or (unfolded as) a set of nodes themselves, and may be either individualized or merged into the same EN. Space has long been modeled in this way, as a graph with changing topologies (Burel & Baudry, 2003; Gaucherel et al., 2012). Conceptually, there is nothing to prevent the merging of the spatial representation in the graph with the more functional EN we have described here: whenever various localities need to be described in the same extensive overall super-graph (Felipe-Lucia et al., 2021), their own EN (but not their corresponding spatial node) may be merged with neighboring ENs too. Hence, each ecosystem may be represented as a changing single hypergraph, including its spatial structure. Although such a single hypernetwork would seem huge, it brings the conceptual advantage to provide a coherent view of any (social-)ecological system, as already mentioned in previous theoretical discussions (Gaucherel, 2019).

## Conclusion

The so-called ecosystem hypernetwork (EN, i.e., the social-ecological interaction hypergraph) fits our need for efficient representation of any complex ecosystem, and provides several additional advantages. It summarizes the internal structure of the system, a kind of synthetic skeleton of the components and processes involved. In this study, we highlighted three points, illustrated with specific Petri net models. Firstly, it is possible to collect all the ecosystem’s interactions into a single EN, regardless of their nature, number and how they intertwine. This proposed representation still shows intentional limitations: for example, in this representation, components never interact directly; here, interactions are always connected to a process. Similarly, no process can impact directly another process without being connected to a component. This single EN is possibly not unique, as ecosystem models are driven by a research question, thus leading to various representations of the same system.

Secondly, the EN is modelled as a hypergraph, which represents non-dyadic interactions more explicitly. Events and components defining the system can be represented as distinct nodes of a bipartite graph, as in Petri nets or, in our case, a qualitative hypernetwork (Fig. 3). Thirdly, the EN representing any ecosystem is dynamic, with a changing topology. We provided an example of a framework (EDEN) computing qualitative ecosystem dynamics based on transitions producing/consuming tokens in the hypergraph. This enables a State-Transition Graph to be computed, collecting all the ecosystem states reachable from the initial state, according to predefined events. This provides a coherent and effective overview of the dynamics of any ecosystem.

## Acknowledgements

We wish to thank Colin Thomas for useful discussions on an early draft of this paper.

## Data, scripts, code, and supplementary information availability

Scripts and code are available online: No data were used in this study; the method sources are archived at https://github.com/fpom/ecco which is the engine as well as the (Jupyter platform) interface of the EDEN framework.

